# Multiple poliovirus-induced organelles suggested by comparison of spatiotemporal dynamics of membranous structures and phosphoinositides

**DOI:** 10.1101/135509

**Authors:** Hyung S. Oh, Sravani Banerjee, David Aponte-Diaz, Jason Aligo, Maria F. Lodeiro, Craig E. Cameron

**Affiliations:** Department of Biochemistry and Molecular Biology, The Pennsylvania State University, University Park, PA 16802

**Author notes:** Correspondence; 814-86-7024; @CameronLabPSU (Twitter). These authors contributed equally to this study (Hyung S. Oh, Sravani Banerjee). Current address: Biologics Toxicology, Janssen Research and Development, LLC, Spring House, PA 19477.

## Abstract

Poliovirus (PV) infection induces membranes with elevated levels of phosphatidylinositol-4-phosphate (PI4P) and invaginated, tubular structures that appear as vesicular clusters in cross section. Here, we characterize PV mutants, termed EG and GG, which exhibit aberrant proteolytic processing of the P3 precursor that can delay the onset of genome replication and/or impair virus assembly. For WT PV, changes to the PI4P pool were observed as early as 30 min post-infection. PI4P remodeling occurred even in the presence of guanidine hydrochloride, a replication inhibitor. Vesicular clusters were not apparent until 3 h post-infection, a time too slow for these structures to be responsible for genome replication. Delays in the onset of genome replication observed for EG and GG PVs were explained completely by the reduced kinetics of virus-induced remodeling of PI4P pools, consistent with PI4P serving as a marker of the genome-replication organelle. Infectious virus produced by GG PV is known to be reduced nearly 5 logs. We show that GG PV was unable to make virus-induced vesicular clusters. Instead, GG PV-infected cells accumulated elongated tubules. Our results are consistent with the existence of distinct organelles for genome-replication and virus assembly. We suggest that the pace of formation and spatiotemporal dynamics of PV induced organelles may be set by the rate of P3 precursor processing to form 3AB and/or 3CD proteins.

**AUTHOR SUMMARY:** All positive-strand RNA viruses replicate their genomes in association with host cell membranes. PV does not just remodel existing membranes, but induces membranes with unique structure and lipid composition. There has been some suggestion that the functions of the PV induced structures observed during infection may not be those that perform genome-replication. This study uses kinetic analysis of virus-induced membrane formation and PI4P induction by two PV mutants to provide evidence for the existence of a virus-induced genome-replication organelle distinct from a second organelle, the absence of which impairs virus assembly. In addition, our studies suggest that formation of both organelles may require participation of viral proteins, 3AB and/or 3CD. Therefore, this study provides a new perspective on the cell biology of PV infection and should inspire a fresh look at picornavirus-induced organelles, their functions and the role of P3 proteins in their formation.

## INTRODUCTION

Positive-strand RNA viruses pose a great threat to public health because of their simplicity and evolvability [1]. Introduction of a modestly-sized, mRNA-sense genome is sufficient to commandeer the cell, establish infection and produce thousands of progeny in a matter of only hours. Among the least understood aspect of positive-strand RNA virus biology is the eclipse phase of the life cycle, the events post entry but prior to exponential amplification of the genome and assembly of infectious virus. We use poliovirus (PV), the type virus in the Enterovirus genus of the *Picornaviridae* family of viruses, as our model system. Establishment of infection by PV likely requires entry of multiple genomes into the cell [2, 3]. This circumstance may be required to permit translation of infecting virion RNA to produce viral protein in sufficient quantity to overwhelm host defenses that might otherwise curtail infection and to hijack host factors essential to initiate a productive infection. It is becoming increasingly clear that picornaviruses have evolved mechanisms to increase multiplicity of infection, for example by leaving cells in vesicles and perhaps entering that way [4] or by attaching to bacterial surface polysaccharides and entering in association with these organisms [5].

Another important function of viral proteins produced during the eclipse phase is establishment of sites of genome-replication. All positive-strand RNA viruses replicate in association with membranes; three general strategies exist [6]. The first strategy is to commandeer an organellar membrane by directing viral proteins to the organelle. Binding of the viral protein will induce invaginations with negative curvature from the perspective of the cytoplasm that have been termed spherules. In response to actions of the same or different viral proteins, the lipid and protein composition of the spherules may be remodeled to reflect the genome-replication needs of the virus. The second strategy uses a two step process. The virus commandeers vesicular and/or organellar membranes, again by directing viral proteins to their membranes. Hijacked membranes are then forced into a new form by remodeling lipid and/or protein composition to create a virus-induced organelle best suited to genome-replication. The third strategy is a blend of the first two. Localize first to an organelle and later induce a virus specific organelle(s), which may be the case for enteroviruses [7].

Since the earliest studies of PV-infected cells using transmission electron microscopy (TEM), it has been well documented that PV infection wreaks havoc on intracellular membranes, for example causing dissolution of the Golgi apparatus (referred to throughout as Golgi), to create membranes of unique benefit to the virus [8-12]. PV induced membranes often appear as clusters of vesicles by TEM. These structures have been shown to contain viral RNA and proteins [13-16]. More recent studies by Ehrenfeld, Belov and colleagues using electron tomography have suggested that PV induced membranes arise from tubules [9]. The thought is that viral or viral-hijacked proteins induce invaginations of positive curvature and promote folding of tubules, which would appear as vesicular clusters in cross section. In the Ehrenfeld and Belov study, a kinetic analysis was performed that provided the first suggestion that the most dramatic PV induced changes to membranes occurred after genome-replication was well underway, if not completed.

Prior to the tomography study, Altan-Bonnet and colleagues showed induction of phosphatidylinositol-4-phosphate (PI4P) by another member of the Enterovirus genus, Coxsackievirus B3 (CVB3) [17]. In uninfected cells, PI4P localizes to the Golgi. During infection, PI4P starts at the Golgi then relocalizes to a site between the endoplasmic reticulum (ER) and Golgi, perhaps the ER-Golgi intermediate compartment (ERGIC). At later times post-infection, PI4P fills the entire perinuclear region of the cell and now localizes with ER markers. During the entire infection period, PI4P localizes with viral RNA and proteins. One conclusion of the study was that PI4P is a marker of the PV induced organelle. When considered in the context of the tomography study, it is possible that the perinuclear PI4P observed for CVB3 reflects the invaginated and folded tubules observed for PV that might not function in genome-replication. Therefore, the earliest stages of PI4P induction and membrane remodeling may actually represent the replication organelle, for which TEM images are not available.

The large polyprotein encoded by the PV genome can be divided into three smaller precursor proteins: P1, P2 and P3 (**Fig. 1A**) [18]. P1 proteins form the capsid. P2 proteins are thought to be the most functionally diverse of the PV proteins as these proteins shut off host cell pathways, derange host cell membranes, function in genome-replication and genome encapsidation, and can even leave their mark on the encapsidated genome by somehow controlling its uncoating [12, 14, 19-30]. P3 proteins are most intimately associated with genome-replication as these proteins bind to the genome and catalyze RNA synthesis [31-37].

**Figure 1.**
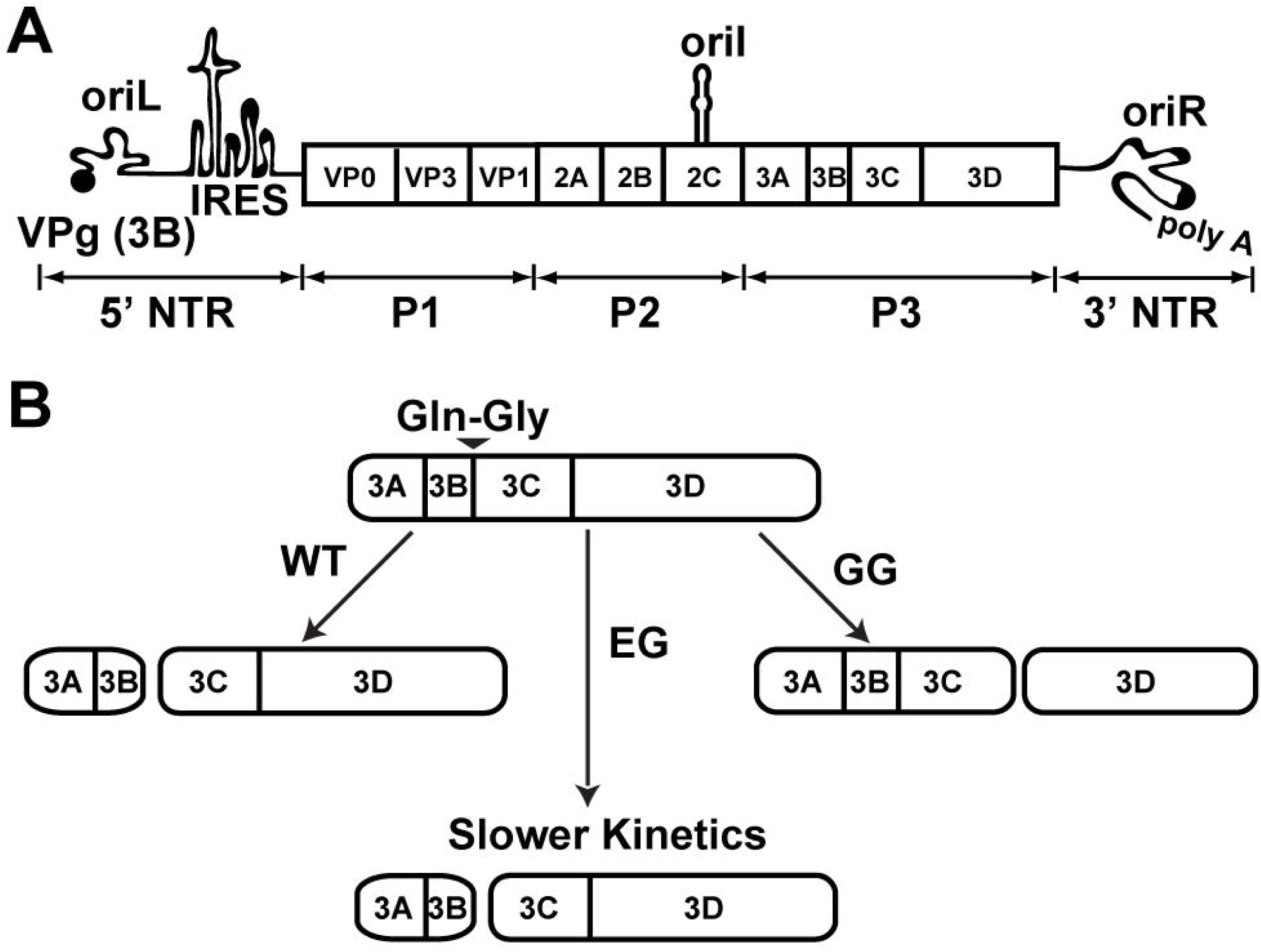
Poliovirus genome organization and P3 polyprotein processing. (**A**) Schematic of the poliovirus genome. The 7.5 kb long genome consists of a 5’ nontranslated region (NTR), an open reading frame, a 3’ NTR and a poly(rA) tail. The 5’ end of the genome is covalently linked to a peptide (VPg) encoded by the 3B gene. The 5’ NTR contains a cis acting replication element (CRE) termed oriL or cloverleaf followed by a type II internal ribosome entry site (IRES). Two additional CREs exist: oriI and oriR, located within the 2C gene and 3’ NTR, respectively. IRES mediated translation yields a single polyprotein comprise of three functional domains of structural (P1) and the non structural (P2 and P3) proteins. (**B**) Processing of the P3 region by WT and mutant PVs. Two pathways, major and minor, exist for P3 processing. Major pathway use for WT PV is shown and produces only 3AB and 3CD because of cleavage at Gln Gly junction between 3B and 3C. EG PV changes the 3B 3C junction to Glu Gly, producing the normal products at a reduced rate. GG PV changes the 3B 3C junction to Gly Gly, which is uncleavable, inducing aberrant processing and producing 3ABC and 3D instead 3AB and 3CD.

PV uses differential cleavage of the polyprotein as one mechanism to expand its proteome. Cleavage of the P3 precursor protein is mediated by the 3C-encoded protease activity. This protease cleaves at Gln-Gly junctions, but the requirements for cleavage extend beyond a single dipeptide [38-40]. Two pathways exist for cleavage of P3 [41, 42]. The major pathway produces only two proteins: 3AB and 3CD (**Fig. 1B**). The minor pathway is more complex. First, P3 is cleaved to produce 3A and 3BCD proteins. 3BCD is then cleaved to produce 3BC and 3D. Finally, 3BC is cleaved to produce 3B (also known as VPg, <u>v</u>irion <u>p</u>rotein linked to the <u>g</u>enome) and 3C.

Previously, we reported two PV mutants in which the cleavage site between 3B and 3C was changed to Gly-Gly (GG PV) [42] or Glu-Gly (EG PV) [42, 43]. For GG PV, the products of the major pathway were 3ABC and 3D, thus ablating the production of both 3AB and 3CD (see GG in **Fig. 1B**). With the exception of 3BC, which cannot be cleaved, products of the minor pathway were essentially unchanged [42]. For EG PV, the products of both pathways were the same as observed for wild type (see EG in **Fig. 1B**). However, the rate of cleavage at the 3BC junction was reduced, causing a reduced rate of accumulation of 3AB and 3CD. Both mutants exhibited what appeared to be a reduced rate of genome-replication. For EG PV, ectopic expression of 3CD alone corrected this phenotype, consistent with the phenotype arising solely from the reduced rate of 3CD production. In addition to the impact on genome yield, GG PV exhibited a near-complete loss of infectious virus. Neither of the GG PV phenotypes could be suppressed by ectopic expression of 3CD.

The extended duration of the eclipse phase observed for EG PV and the defect to virus assembly observed for GG PV provided a unique opportunity to expand our understanding of these steps of the virus lifecycle at the molecular and ultrastructural levels. We find that there is a clear temporal ordering of PI4P induction and its initial induction and limited redistribution exhibit the expected kinetics for an organelle involved in genome-replication. During the early stages of PI4P remodeling, Golgi fragments and ERGIC expands. It is only after genome-replication is near completion that the perinuclear region of the cell becomes filled with vesicular clusters. At the same stage of genome-replication that vesicular clusters predominate for WT PV, tubules predominate for GG PV. The inability of GG PV to assemble infectious virus is consistent with the possibility that vesicular clusters contribute substantively to steps after genome-replication on path to production of infectious virus. We propose that two organelles are induced during PV infection that serve distinct functions during the PV lifecycle and that P3-encoded proteins contribute to formation and function of both.

## RESULTS

### Delayed release of 3AB and 3CD proteins from the P3 precursor protein delays onset of genome-replication

Our previous studies of EG PV showed that its genome-replication defect could be complemented in trans by expression of 3CD [43]. This observation was unexpected because numerous studies have shown that genome-replication defects can only be complemented in trans by expression of P3 or P2-P3 polyproteins, if at all [44-46]. Our study used a subgenomic replicon in which P1-coding sequence had been replaced with firefly luciferase-coding sequence, and replication was monitored indirectly by measuring luciferase activity. It was possible that measuring luciferase activity instead of RNA confounded our interpretation of the experiment.

To test this possibility, we transfected WT or EG PV subgenomic replicon RNA into HeLa cells. We monitored RNA synthesis either by measuring luciferase activity (**Fig. 2A**) or by measuring positive-sense RNA using Northern blotting (**Figs. 2B** and **2C**). We observed three discernible phases of luciferase activity for WT: 0-2 h, 2-5 h, and 5-10 h; and for EG: 0-3 h, 3-7 h and 7-10 h (**Fig. 2A**). In contrast, we observed only one phase of RNA accumulation for both (**Fig. 2B**). WT RNA accumulation began between 1-2 h post-transfection; EG RNA accumulation began between 3-4 h post-transfection (**Fig. 2B**). We conclude that a delay of two hours in the onset of replicon RNA replication can be obscured by monitoring luciferase activity. Luciferase activity concealed the delay. Translation of transfected replicon RNA produced luciferase activity for at least two hours in the presence of guanidine hydrochloride (GuHCl), a replication inhibitor, and therefore without the need for replicon RNA replication (see WT +GuHCl in **Fig. 2A**). These results support the hypothesis that delayed kinetics of P3 precursor processing to form 3AB and 3CD in cells causes a delay in a step preceding the onset of RNA synthesis.

**Figure 2.**
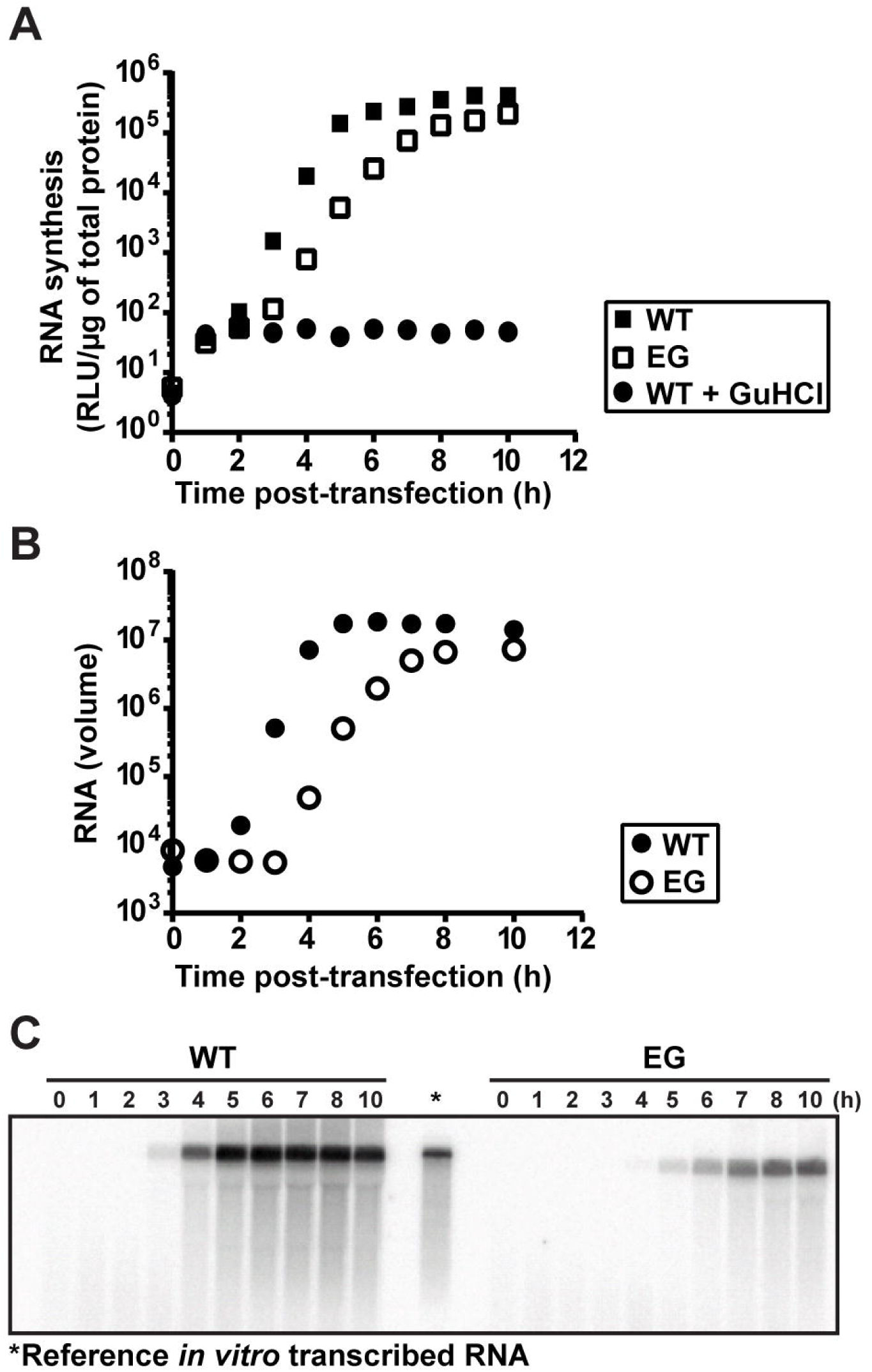
EG PV exhibits a delay in the onset of RNA synthesis. (**A**) Kinetics of replication of subgenomic replicon by WT (▪) and EG (▫) monitored indirectly by luciferase activity. HeLa cells were transfected with *in vitro* transcribed replicon RNA, placed at 37 °C and luciferase activity (RLU/μg) measured at the indicated times post transfection. A control for translation of transfected replicon RNA was performed by performing the experiment above in the presence of 3 Mm GuHCl (•). (**B**) Kinetics of replication of subgenomic replicon by WT (•) and EG (○) monitored by Northern blotting. HeLa cells were transfected with *in vitro* transcribed replicon RNA, placed at 37 °C, and at the indicated times post transfection, cells were harvested for total RNA isolation. Total RNA was separated on a 0.6% agarose gel containing 0.8 M formaldehyde, transferred to nylon membrane and hybridized with a ^32^P labeled DNA probe. (**C**) Image of a representative blot visualized by phosphorimaging. *In vitro* transcribed RNA (**\***) is shown as reference.

### Kinetics of formation of PV-induced vesicular clusters are inconsistent with these structures serving as the sites of genome-replication

A hallmark of PV-infected cells as observed by transmission electron microscopy (TEM) is the presence of vesicular clusters [9, 12]. These vesicular clusters are also referred to as replication complexes or replication organelles. Given the accepted role of these vesicular clusters in genome-replication, we reasoned that formation of these structures should occur either prior to the onset of genome-replication or concomitant with genome-replication. Formation and/or function of these structures could therefore represent the pre-genome-replication step perturbed for EG PV.

We infected HeLa cells with EG PV and then at various times post-infection processed samples to measure genomic RNA by Northern blotting (**Figs. 3A** and **3B**), to measure infectious virus by plaque assay (**Fig. 3A**) and to visualize formation of the vesicular clusters by TEM (**Fig. 3C**). The exponential phase of genome-replication occurred between 2 h and 6 h post-infection (**Figs. 3A** and **3B**). The exponential phase of infectious virus production occurred between 4 h and 8 h post-infection (**Fig. 3A**). Vesicular clusters became visible at 4 h post-infection and continued to increase in abundance and/or change in organization throughout the 7 h time course (**Fig. 3C**). Formation of vesicular clusters neither preceded genome-replication nor occurred concomitant with genome-replication as expected for an organelle responsible for genome-replication. In contrast, formation of vesicular clusters preceded infectious virus production.

**Figure 3.**
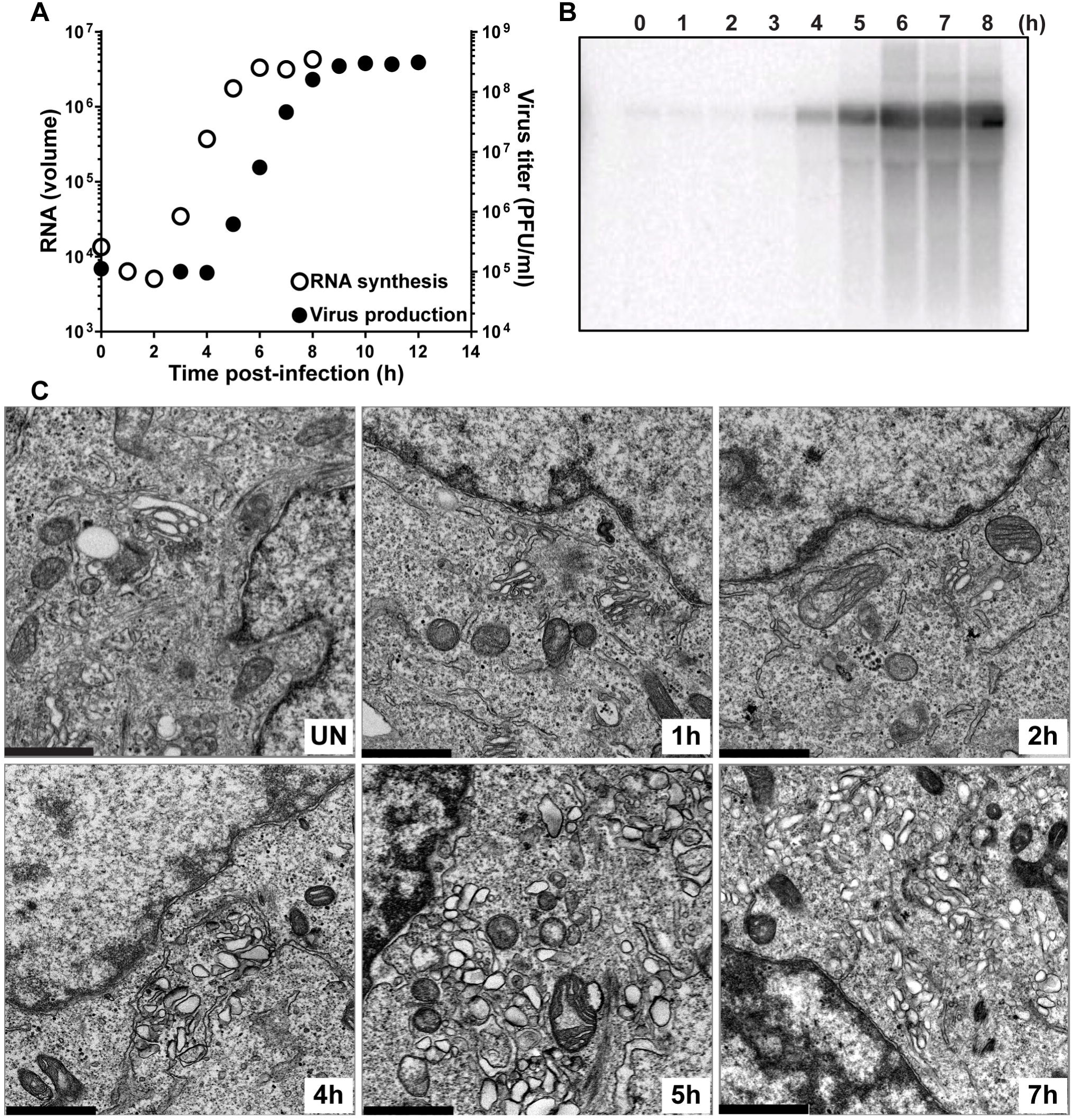
Kinetics of genome-replication precedes the kinetics of vesicular cluster formation for EG PV. (**A**) Kinetics of RNA synthesis (○) and virus production (•) by EG PV. HeLa cells were infected with EG PV at an MOI of 10, placed at 37 °C, and at the indicated times post-infection, total RNA was isolated and subjected to either Northern blotting or assayed for virus production by standard plaque assay. (**B**) Image of a representative blot visualized by phosphorimaging. **(C)** Kinetics of formation of virus-induced vesicular clusters by transmission electron microscope (TEM). HeLa cells were infected with EG PV at MOI of 10, placed at 37 °C, and at the indicated times post-infection, infected cells were fixed and visualized by TEM, bar = 1 μm.

The delay in induction of the vesicular clusters by EG PV provided a unique opportunity to monitor the early steps in formation of these structures. In uninfected cells, it was very easy to observe mitochondria and Golgi. However, closer inspection was required to observe the tubules of the endoplasmic reticulum (ER) (uninfected in **Fig. 3C**). By 2 h post-infection, the density of mitochondria had diminished (2 h in **Fig. 3C**). Golgi stacks were fragmented and swollen, and the tubules of the ER and/or ER-Golgi intermediate compartment (ERGIC) were swollen and easier to visualize (2 h in **Fig. 3C**). At 3 h post-infection, much of the Golgi had disappeared (3 h in **Fig. 3C**). The area between the remaining Golgi and the now very swollen ER was filled with very small vesicles that may be the ever expanding ERGIC caused by the loss of the Golgi (3 h in **Fig. 3C**) [17]. At 4 h post-infection, vesicular clusters were first observed (4 h in **Fig. 3C**). These vesicular clusters appear to be encased by the swollen tubules of the ER (4 h in **Fig. 3C**).

In order to make certain that our observations above were not unique to the mutant virus, we repeated the experiment using WT PV (**Fig. 4**). The exponential phase of genome-replication occurred between 1 h and 5 h post-infection (**Figs. 4A** and **4B**). The exponential phase of infectious virus production occurred between 3 h and 7 h post-infection (**Fig. 4A**). Vesicular clusters were now clearly visible at 3 h post-infection and again continued to increase in abundance or change in organization throughout the 7 h time course (**Fig. 4C**). Many of the early steps described for EG PV could not be observed for wild type because they occurred so soon post-infection that we could not process the cells fast enough to capture images of these steps by TEM.

**Figure 4.**
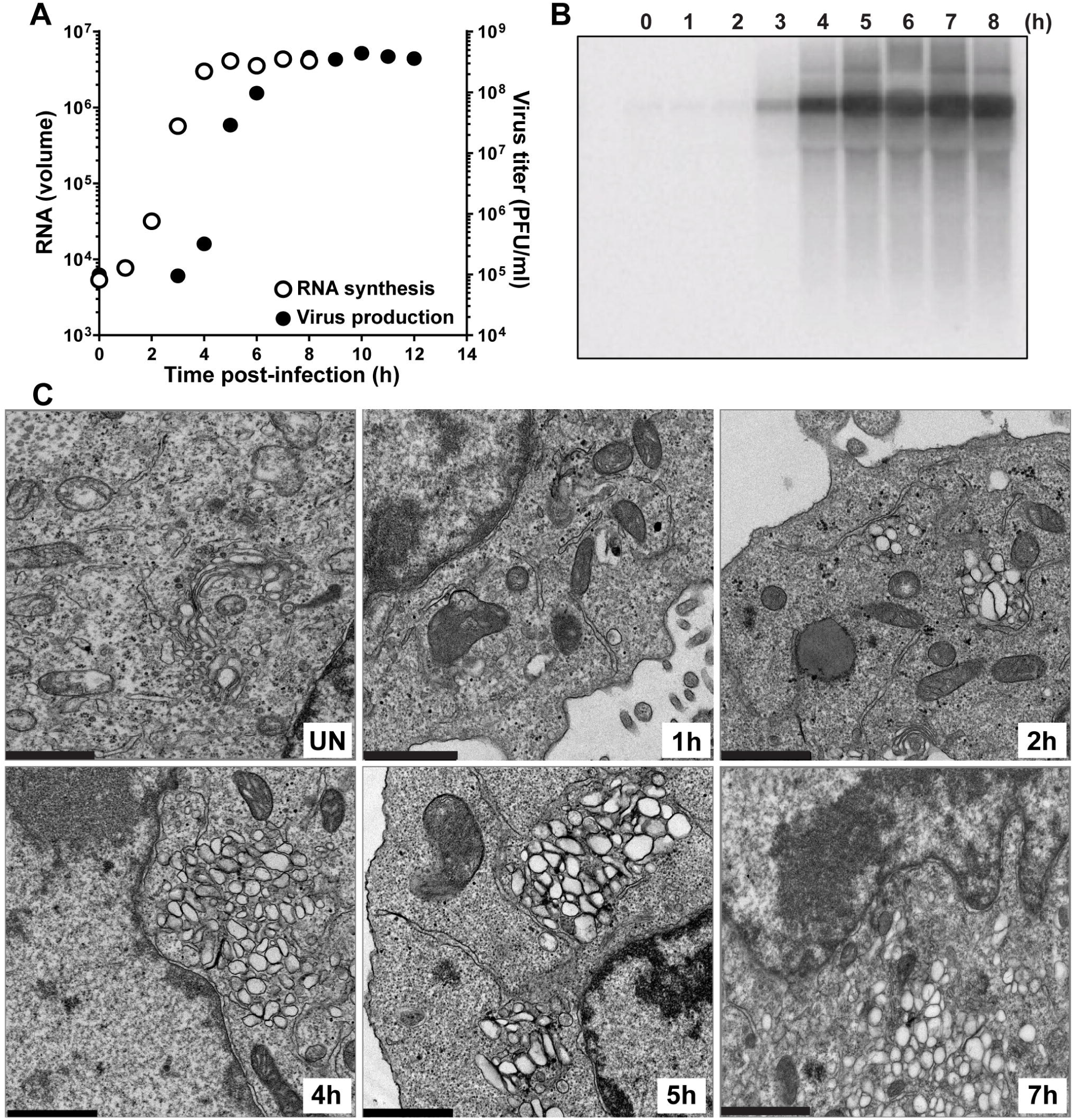
Kinetics of genome-replication precedes the kinetics of vesicular cluster formation for WT PV. (**A**) Kinetics of RNA synthesis (○) and virus production (•) by WT PV. HeLa cells were infected with WT PV at an MOI of 10, placed at 37 °C, and at the indicated times post-infection, total RNA was isolated and subjected to either Northern blotting or assayed for virus production by standard plaque assay. (**B**) Image of a representative blot visualized by phosphorimaging. (**C**) Kinetics of formation of WT PV-induced vesicular cluster formation was visualized by TEM. HeLa cells were infected with WT PV at an MOI of 10, placed at 37 °C, and at the indicated times post-infection, the infected cells were fixed and visualized by TEM, bar = 1 μm.

In addition to the temporal difference in the formation of the vesicular clusters observed for EG PV and WT PV, differences also existed in the overall size, interaction and localization of these structures (compare 5 h for EG PV in **Fig. 3C** to 4 h for WT PV in **Fig. 4C**). Relative to EG PV, vesicular clusters induced by WT PV appeared larger, clustered more tightly and were far more constrained to the perinuclear region of the cell.

Both experiments were consistent with the surprising finding that the kinetics of formation of the PV-induced vesicular clusters are inconsistent with these structures serving as the genome-replication organelle. These structures became easily identifiable two hours after genome-replication began (4 h for EG PV in **Fig. 3C**; 3 h for WT PV in **Fig. 4C**). Interestingly, and perhaps coincidentally, this was the time after genome-replication began that we were able to detect infectious virus by plaque assay (**Figs. 3B** and **4B**).

### Kinetics of PI4P induction are consistent with this phosphoinositide demarcating the sites of genome-replication

Coxsackievirus B3 (CVB3) infection induces PI4P and this phosphoinositide is a component of a virus-induced organelle [17]. PI4P co-localizes with viral non-structural proteins and RNA [17]. Whether or not PI4P localizes to the virus-induced vesicular clusters is not known. Given our results, it is possible that the virus-induced PI4P localizes to membranes other than those observed by TEM well after genome-replication has begun.

Although it is assumed that PV infection induces PI4P based on the data for CVB3, PI4P induction by PV has never been reported. We infected HeLa cells with WT PV or EG PV and measured PI4P as a function of time post-infection by immunofluorescence microscopy (IFM) (**Fig. 5**). We used an antibody specific for PI4P to detect this phosphoinositide.

**Figure 5.**
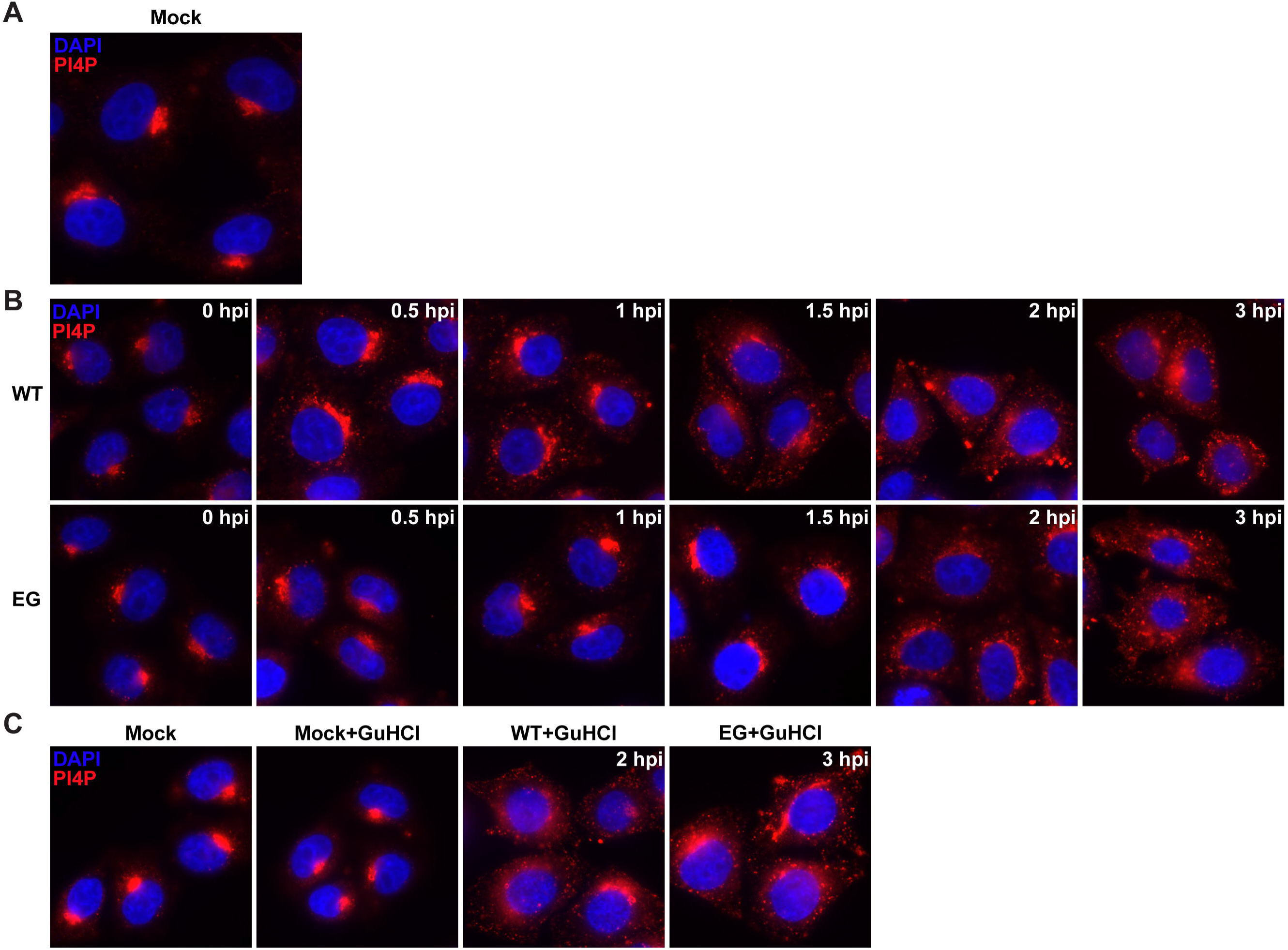
EG PV exhibits delayed induction and redistribution of PI4P. (A) Immunostaining of PI4P in mock infected HeLa cells. **(B)** Time course of PI4P staining in HeLa cells infected with WT or EG PV. HeLa cells were infected with WT or EG virus at an MOI of 10, fixed at indicated times post-infection, and immunostained for PI4P. **(C)** Impact of GuHCl on PI4P induction by WT and EG PVs. HeLa cells were either incubated with PBS or infected with WT or EG virus (MOI 10) in presence of 3 mM GuHCl and immunostained. In all cases, PI4P was stained using anti PI4P antibody (red) and nuclei were stained with DAPI (blue).

It is well known that most PI4P in the cell localizes to the Golgi, giving rise to very nice staining with anti PI4P antibody (mock in **Fig. 5A**) [17]. For WT PV, induction and redistribution of Golgi localized PI4P was evident as early as 30 min post-infection WT in **Fig. 5B**). Both redistribution and induction continued to 3 h post-infection, the last time point evaluated (WT in **Fig. 5B**). Observations with EG PV were essentially identical, except for a one hour delay of the changes relative to wild type (EG in **Fig. 5B**). The kinetics of induction and redistribution of PI4P were consistent with a structure responsible for genome-replication. Given the TEM experiments reported above (**Figs. 3** and **4**), it is now clear that the PI4P containing membranes observed by IFM prior to two hours post-infection (**Fig. 5B**) must be distinct from the large vesicular clusters that appear after genome-replication (**Figs. 3** and **4**).

PI4P induction and redistribution during WT PV infection occurred as fast as we could reliably process our cells for imaging. This observation suggested to us that viral proteins made by translation of infecting viral RNA might be sufficient to initiate PI4P remodeling. We infected cells with WT or EG PV in the presence of the genome-replication inhibitor, GuHCl. Under these conditions, the only event that can occur is translation of the infecting genomes. PI4P induction and redistribution occurred in the presence of GuHCl for both WT (WT+GuHCl in **Fig. 5C**) EG (EG+GuHCl in **Fig. 5C**) PVs. GuHCl did not have an impact on PI4P localization in the absence of infection (Mock+GuHCl in **Fig. 5C**).

### PV-induced vesicular clusters as sites of virus assembly, an assembly organelle?

The studies reported above suggested that PI4P was a marker for the PV genome-replication organelle. These studies also suggested that PI4P-containing structures observed early are distinct from PV-induced vesicular clusters. The question now was: what is the function of the vesicular clusters? One possibility was that the vesicular clusters are structures required for virus assembly, an assembly organelle, as suggested previously by Belov and Ehrenfeld [9]. Another possibility was that the vesicular clusters are structures that represent accumulated, unused and unwanted viral proteins and nucleic acids, a viral trash compacter.

GG PV replicated to within one log of wild type but exhibited a near 5-log reduction in the amount of infectious virus produced [42]. We reasoned that this mutant might help us identify the function of the PV-induced vesicular clusters. If the vesicular clusters contribute to virus assembly, then these structures might be perturbed in cells infected with the assembly-defective GG PV. Conversely, if the vesicular clusters function as a means to consolidate viral waste, then these structures might appear unchanged in cells infected with GG PV.

We transfected HeLa cells simultaneously with two subgenomic replicons, one expressing luciferase and the other expressing EGFP. Transfected cells were then grown in suspension culture and harvested hourly to monitor luciferase activity and EGFP fluorescence. Luciferase specific activity, measured as relative light units per microgram of protein in the extract, was plotted as a function of time for WT and GG replicons (**Fig. 6A**). We were particularly interested in 5 h post transfection for WT as vesicular clusters reached their maximum level at that time (**Fig. 4C**). At 5 h post transfection of the WT replicon combination, the luciferase specific activity was approximately 10^4^ RLU/μg (**Fig. 6A**). We observed the equivalent luciferase specific activity at 14 h post transfection of the GG replicon combination (**Fig. 6A**). Based on this result, we concluded that 5 h and 14 h post transfection represented equivalent extents of the infection process for WT and GG PVs, respectively.

**Figure 6.**
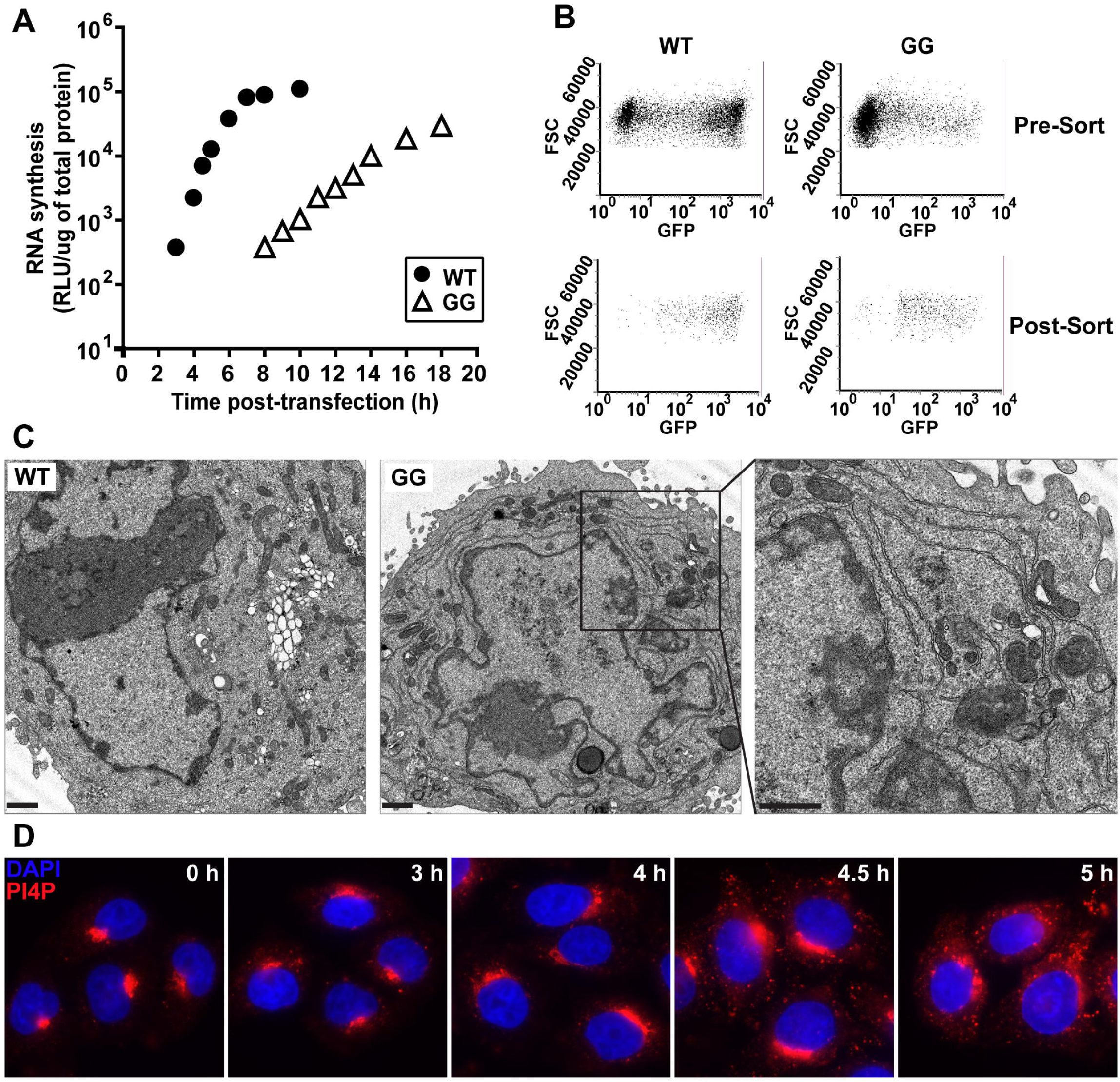
GG PV induces and redistributes PI4P in spite of impaired formation of vesicular clusters. (**A**) Kinetics of RNA synthesis by WT (•) and GG (?) subgenomic replicon RNA. HeLa cells were co transfected with two different replicon RNAs, luciferase replicon (2 μg) and EGFP replicon (4 μg), placed at 34 °C and at the indicated times post transfection, luciferase activity was measured. (**B**) Cell sorting to isolate PV replicon positive cells. WT replicon RNA transfected cells were 61% positive in pre sort cells (top left) and 98% positive in post sort cells (bottom left). GG replicon RNA transfected cells were 18% positive in pre sort cells (top right) and 94% positive in post sort cells (bottom right). (**C**) WT and GG PV-induced membranes visualized by TEM. HeLa cells were transfected with either WT or GG replicon RNA, placed at 34 °C for 5 h or 14 h, respectively, at which time cells were fixed and visualized by TEM. Bar =1 μm. **(D)** Kinetics of PI4P induction and redistribution by the GG PV subgenomic replicon. HeLa cells were transfected with replicon RNA expressing EGFP and samples were fixed at the indicated time post transfection and subjected to IFM using anti PI4P antibody (red) and nuclei were stained with DAPI (blue).

We used fluorescence activated cell sorting (FACS) to isolate transfected cells based on EGFP expression. Cells replicating WT or GG EGFP replicon RNA before and after sorting are shown in **Fig. 6B**. We used the EGFP expressing, sorted cells for TEM (**Fig. 6C**). Vesicular clusters were readily apparent in images of WT replicon-transfected cells, and these clusters were of the same overall appearance as observed using infected cells that did not require sorting (**Fig. 4C**). In contrast, vesicular clusters were few or absent in images of GG replicon-transfected cells (**Fig. 6C**). Instead, we observed a tubular/reticular network (see enlargement of panel GG in **Fig 6C**).

These results support the hypothesis that vesicular clusters function as an assembly organelle, and the absence of these structures in GG PV contribute to the defect in virus production. We propose that the tubular/reticular network is an intermediate on the pathway to formation of the vesicular clusters. These structures were apparent between 2 h and 4 h post-infection for EG PV (**Fig. 3C**) but occurred too early to be observed for WT PV (**Fig. 4C**). The tubules appeared to fold in on themselves, creating the vesicular clusters (see for example 4 hpi for EG PV in **Fig. 3C** or 3 hpi for WT PV in **Fig. 4C**). Formation of tubules and their transformation into a net-like array of vesicles has been observed before [9]. Additional studies will also be required to further clarify the relationship between the tubular/reticular network in GG PV and the vesicular clusters in EG PV and WT PV.

Finally, we assessed formation of the PI4P-marked structures by GG PV RNA. We transfected HeLa cells with GG PV EGFP-expressing, subgenomic replicon RNA and used IFM to monitor the spatiotemporal dynamics of PI4P. Redistribution and induction of PI4P occurred substantially slower than observed for EG PV or WT PV (compare **Fig. 6D** to **Fig. 5**). Nevertheless, temporal dynamics of PI4P explains the 8 h or so pre-genome-replication phase observed previously for GG PV [42, 43]. These data are consistent with PI4P demarcating the replication organelle and a requirement for induction of this structure prior to the onset of genome-replication.

### High multiplicity of infection exaggerates spatiotemporal dynamics of vesicular clusters

The spatiotemporal dynamics of WT PV-induced vesicular clusters reported here were different than observed previously by others [9, 11, 16]. In particular, formation of the clusters was slower, and the density of the clusters was lower (**Fig. 4C**) than previously reported. In these earlier experiments, multiplicity of infection ranged from 30-50, but this study used 10. Because translation of the infecting genomes is sufficient to begin the remodeling process, it was possible that elevated levels of protein produced at the higher multiplicities of infection might impact the size of the clusters. To address this possibility, we infected HeLa cells with WT PV at a multiplicity of 10, 50 or 100 and processed the infected cells for TEM at 3 h post-infection. This time point was chosen because of the modest density of the vesicular clusters observed at a multiplicity of infection of 10 (**Fig. 4C**) and should make any observed differences more noticeable. As shown in **Fig. 7**, the size of the vesicular cluster at the level of each “vesicle” and in terms of the area of the perinuclear region occupied by the vesicular clusters scaled directly with the multiplicity of infection. Therefore, it is imperative to consider multiplicity of infection when comparing micrographs.

**Figure 7.**
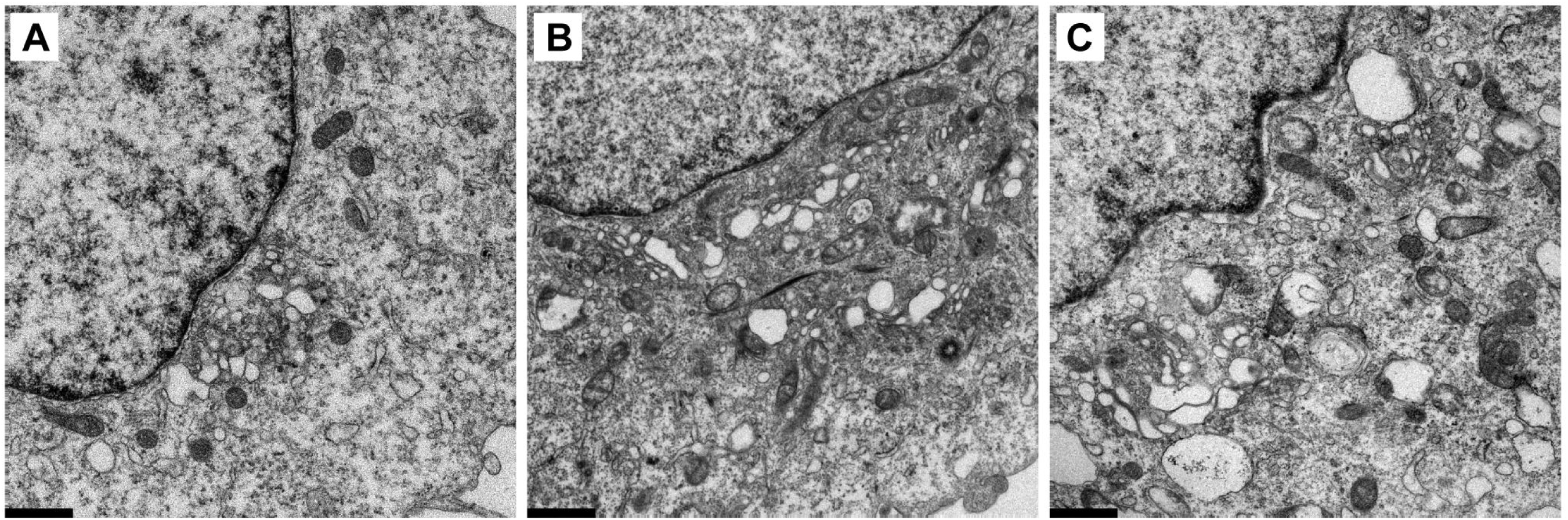
Multiplicity of infection exaggerates size and quantity of WT PV-induced vesicular clusters. Transmission electron micrographs (TEM) of HeLa cells infected with WT PV at an MOI of 10 **(A),** 50 **(B)** or 100 **(C)**. HeLa cells were infected with WT PV expressing EGFP at an MOI of 10, 50 or 100 by incubating the monolayer with WT virus for 30 min at room temperature, followed by removal of the virus and washing of the monolayer with PBS. Prewarmed media was then added to the monolayer and incubated at 37 °C. Samples were fixed 3 h post-infection and visualized by TEM. Bar = 1 μm.

## DISCUSSION

This study was inspired by our observation that ectopic expression of PV 3CD protein was sufficient to complement a genome-replication defect of the EG PV mutant, which exhibited a delay in processing of the P3 precursor protein [43]. The rule of thumb for PV and other picornaviruses is that non structural proteins used to replicate a genome are produced in cis—that is, by that genome or by one in the same replication complex [44-46]. However, it became clear that the genome-replication defect was actually a defect at steps preceding genome-replication (**Fig. 2B**). The confusion was caused by the multiphasic kinetics of reporter activity produced by a subgenomic replicon (**Fig. 2A**). The first phase of activity requires only translation of the transfected replicon. The second phase requires replicon replication. The third phase appears to be translation of replicated replicon RNA that would normally be packaged if capsid proteins had been present.

Once we knew that EG PV impaired steps leading up to genome-replication, we turned our focus to formation of the “replication organelle.” Since the first TEM images of PV-infected cells were published [8], multiple reports have shown that a hallmark of PV infection is the induction of membranous structures [9-12]. These structures have many names: replication complexes, vesicular clusters, tubules and replication organelle. Most of the early studies searching for viral factors contributing to formation of PV-induced membranes implicated P2 proteins in this process [14, 15]. The most convincing experiment showed that expression of the 2BC precursor proteins was sufficient to induce membranes with the same buoyant density and appearance by TEM as those observed during PV infection [20, 21]. As a result, it was unclear how processing of the P3 precursor to form 3AB and 3CD contributed to formation of the replication organelle. However, because formation of this structure should precede genome-replication, it was possible that this step could be complemented in trans. 3A is known to inhibit guanine nucleotide exchange factor (GEF), GBF1, which prevents trafficking between ER and Golgi and thereby causes dissolution of the Golgi [47-52]. 3AB might also have this activity. Inhibition of ER-to-Golgi trafficking should enlarge the ER-Golgi intermediate compartment, and this compartment has been implicated in genome-replication [17, 51]. 3AB is also known to promote invagination of membranes, an activity that could contribute to formation of vesicular clusters [53]. 3CD, on the other hand, has not been shown to perturb membrane form or function in cells. However, in cell free extracts, the presence of 3CD promotes recruitment of ADP ribosylation factors (ARFs) and GEFs to membranes, which could modulate vesicular trafficking and/or fusion [54, 55].

Our examination of the kinetics of formation of the vesicular clusters by EG PV relative to the kinetics of genome-replication was eye-opening. The kinetics of these two processes did not have the expected behavior---that is, the site of RNA synthesis forming prior to RNA synthesis (**Fig. 3**). This unexpected kinetic behavior was not an oddity of the mutant as WT PV also exhibited this behavior (**Fig. 4**). Our conclusion was that the vesicular clusters must not be the replication organelle. Belov and Ehrenfeld floated this possibility several years ago [9]. In reviewing the literature on formation of PV-induced vesicles, it became clear that the TEM images of others showed faster formation of vesicular clusters in spite of the fact that kinetics of genome-replication and/or production of infectious virus were identical [9, 11, 16]. The one difference between our study and those of others was the multiplicity of infection used. We used an MOI of 10 pfu/cell; others used an MOI of 30-50 pfu/cell. When we increased the MOI to these higher levels, we observed faster kinetics of formation of the vesicular clusters (**Fig. 7**). Under these conditions, the vesicular clusters were also substantially larger (**Fig. 7**). These observations suggest that the kinetics of vesicular cluster formation can be modulated by multiplicity of infection without a concomitant increase in the kinetics of genome-replication or virus assembly, thus creating some ambiguity to the temporal ordering of formation of the vesicular clusters relative to replication of the genome.

Altan-Bonnet and colleagues showed that phosphatidylinositol-4-phosphate (PI4P) is induced by Coxsackievirus B3 infection and redistributes from the Golgi to sites thought to be involved in genome-replication [17]. Therefore, we monitored the spatiotemporal dynamics of PI4P during PV infection. We found that the spatiotemporal dynamics of the PI4P pool was completely consistent with PI4P demarcating the replication organelle as induction and redistribution occurred well before the onset of genome-replication (**Fig. 5A**). Moreover, spatiotemporal dynamics of PI4P was delayed for EG PV (**Fig. 5B**). Unexpectedly, PI4P induction and redistribution did not require any genome-replication as these events occurred in the presence of a replication inhibitor. To the best of our knowledge, this observation is among the first to link translation of infecting viral genomes alone to events essential for genome-replication. Under these conditions, the amount of viral proteins produced cannot be detected readily by Western blotting or immunofluorescence [42, 43]. Therefore, very little of one or more viral proteins is sufficient to hijack cellular factors needed for PI4P induction and redistribution.

We propose that 3AB and/or 3CD contribute to PI4P induction and/or redistribution. If this is the case, then delays in P3 processing would be expected to have an impact on PI4P dynamics. Our initial characterization of EG PV showed that its genome-replication phenotype could be complemented fully by ectopic expression of 3CD [43]. It is therefore possible that 3CD or 3CD-containing precursor alone contribute to PI4P dynamics. Interestingly, synthesis of PI4P by the beta isoform of the class III phosphatidylinositol-4-kinase (PI4KIIIß), the isoform thought to be responsible for induction of PI4P by enteroviruses [17], is known to require Arf1 and GBF1 [17]. Perhaps the ability of 3CD to induce translocation of Arf1 and GBF1 from the cytoplasm to membranes may be related to its role in PI4P induction suggested here [54, 55].

The delay in formation of the replication organelle by EG PV created a window into the early events of this process. It was possible to visualize Golgi fragmentation, proliferation of vesicles in the area between ER and Golgi, and formation of short, thick tubules (**Fig. 3**). These events were suggested by Altan-Bonnet and colleagues based on Golgi, ERGIC and ER marker co-localization with PI4P and CVB3 non structural proteins and/or RNA [17]. Together, these data make a compelling case for these localized membranous structures including and between ER and Golgi representing the replication organelle.

Clues into the formation and function of PV-induced vesicular clusters were obtained from studies of GG PV. This mutant took even longer to initiate genome-replication than EG PV, and this delay was attributable to an even longer delay in PI4P induction and redistribution (**Fig. 6**). GG PV does not produce any 3AB or 3CD, which is likely the cause of this phenotype [42, 43]. This mutant clearly establishes a role for P3 proteins in formation of the replication organelle. However, TEM of GG PV-infected cells revealed a cell full of tubules at times in which genome-replication of GG PV RNA was occurring exponentially (**Fig. 6**). Hints of tubulation were evident in micrographs of EG and WT PVs (**Figs. 3 and 4**), but tubules never really accumulated. We propose that the tubules observed in GG PV-infected cells are an intermediate on path to formation of the vesicular clusters. Positive invagination of the tubules and interactions between invaginated membranes should produce vesicular clusters, as seen in cross section by TEM. As to function of the vesicular clusters, GG PV was unable to produce significant titers of infectious virus [42, 43]. We propose that formation of vesicular clusters requires 3AB and/or 3CD and facilitates virus assembly. It is known that 3CD promotes cell-free synthesis of infectious PV [56, 57]. It is possible that these membranes serve as an assembly organelle.

There was a time when poliovirus non-structural proteins were considered in two categories: those that had an impact on interactions with the host; and those that had an impact on genome-replication. Emphasis was often placed on shut off of host protein synthesis, permeabilization of cellular membranes and formation of vesicles by P2-encoded proteins. P3-encoded proteins bind to the viral genome, produce the peptide/precursor used to initiate genome-replication and function to replicate the genome. This study highlights the almost unlimited functional capacity of PV non-structural proteins by now implicating P3 proteins in events both before and after genome-replication. Understanding how these proteins contribute to these events requires further study. It is likely that the findings reported here for PV will apply to other picornaviruses as well.

## MATERIALS AND METHODS

### Materials

Cell culture media and supplements were from Invitrogen Life Technologies (Gibco); Difco-NZCYM for bacterial growth was from BD Biosciences; 5X Cell Culture Lysis Reagent was purchased from Promega; primary antibody against PI4P was purchased from Echelon biosciences; Alexafluor-594-IgM (goat anti-mouse) was from ThermoFisher Scientific; 16% formaldehyde, methanol-free, ultrapure was purchased from polysciences; Digitonin was from Sigma-Aldrich; Bovine Serum Albumin (BSA)-Fraction V, heat shock treated was from Fisher Bioreagents; Vectashield Antifade Mounting Medium with DAPI was from Vector Laboratories; nitrocellulose membrane was from GE Healthcare; TransIT-mRNA transfection kit was purchased from Mirus Bio LLC.

### Construction of 3B-3C Gly-Gly mutant EGFP subgenomic replicon

Enhanced green fluorescence protein (EGFP) integrated subgenomic replicon plasmid, pREGFP-, was generated by replacing luciferase gene in pRLuc [42] with EGFP. PCR was used to amplify the EGFP region using the oligonucleotides NotI-EGFP-for (5’-AAT TCG GAG CGG CCG CTG TGA GCA AGG GCG AGG AGC-3’) and EGFP-XhoI-rev (5’-GTC AGA TCC TCG AGC TTG TAC AGC TCG TCC ATG-3’) and pEGFP as a template. The PCR product replaced luciferase gene in pRLuc-GG using NotI and XhoI sites to obtain pREGFP-GG subgenomic replicon. Sequencing at the Penn State Nucleic Acid facility was used to confirm all clones.

### RNA transcription

Linearization and RNA transcription were performed as described previously [42]. Briefly, the pMo-, pRLuc- and pREGFP- plasmids were linearized with EcoRI or ApaI and purified with Qiaex II bead suspension (Qiagen) following manufacture’s protocol. RNA was then transcribed from the linearized plasmid DNAs in a 20 μL reaction containing 350 mM HEPES, pH 7.5, 32 mM magnesium acetate, 40 mM dithiothreitol (DTT), 2 mM spermidine, 28 mM nucleoside triphosphates (NTPs), 0.025 μg/μL linearized DNA, and 0.025 μg/μL T7 RNA polymerase. The reaction mixture was incubated for 3 h at 37°C and magnesium pyrophosphate was removed by centrifugation for 2 min. The supernatant was transferred to a new tube and subjected to RQ1 DNase treatment (Promega) for 30 min at 37°C. RNA quality was then verified by 0.8% agarose gel electrophoresis. The RNA concentration was determined by comparing pre-known purified RNA side-by-side on an agarose gel.

### Subgenomic luciferase replicon assay

Subgenomic luciferase assays were performed as described previously [42] with the following modifications. Subgenomic replicon RNA (5 μg of *in vitro* transcribed RNA) was electroporated into HeLa cells (American Type Culture Collection). The cells were incubated in normal growth media (DMEM/F12 supplemented with 10% fetal bovine serum, 1% penicillin/streptomycin, 5 mL/1 X 10^6^ cells) and 1 X 10^5^ cells were harvested and lysed using 100 μL of 1X cell culture lysis reagent (CCLR) at an indicated time post electroporation. Luciferase activity was measured by adding an equal volume of firefly luciferase assay substrate to cell lysate and the reaction mixture was applied to a Junior LB 9509 luminometer (Berthold Technologies) to read relative light units (RLU) for 10 s. Relative light units (RLU) were then normalized based on the total protein concentration determined by Biorad protein assay reagent (BioRad) for each sample. GuHCl when used, the final concentration was 3 mM and the inhibitor was kept on cells during transfection and after, for the entire incubation period.

### Total RNA preparation

One day prior to infection, 4 X 10^6^ HeLa cells were seeded in a 100 mm dish. On the day of infection, the prepared HeLa monolayers were infected at an MOI 10 with WT or EG PV for 30 min at room temperature. Infected cells were washed once with PBS to remove unattached virus and then incubated in normal growth media (DMEM/F12 supplemented with 10% fetal bovine serum and 1% penicillin/streptomycin) at 37°C. At the indicated times post-infection, cells were lysed with 1 mL of Trizol (Invitrogen Life Technologies) and total RNAs were purified by following manufacture’s protocol. In order to purify subgenomic replicon RNA containing total RNA, HeLa cells were transfected with subgenomic RNA as described above and 1.2 X 10^6^ HeLa cells were lysed with 1 mL Trizol. The RNA quantity was determined using NanoDrop 1000 (ThermoFisher Scientific) and the quality was evaluated by using 0.8% agarose gel.

### Northern-Blot analysis of RNA synthesis

Northern-Blot analysis was performed as described previously [43]. Briefly, the RNA was separated on a 0.6% denaturing agarose gel (0.8 M formaldehyde in 1X MOPS) by running at 120V for 2 h. The gel was washed in water for 30 min twice followed by soaking in 20X SSC (Saline Sodium Citrate) buffer for 30 min. RNA was transferred to nylon membrane (Hybond XL, GE Healthcare) using capillary blotting with 20X SSC for 16 h at room temperature. RNA was crosslinked to membrane using UV Crosslinker (Stratalinker 2400, Stratagene). The membrane was washed twice with wash buffer (1X SSC and 0.1% SDS) at 65°C for 30 min each and prehybridized in 100 mL of modified church buffer (0.5 M sodium phosphate, pH 7.2, 7% SDS, and 1 mM EDTA) for 4 h at 65°C. Hybridization was performed in a modified church buffer for 16 h at 65°C. The membrane was washed with wash buffer (1X SSC and 0.1% SDS) for 20 min at 65°C twice and once at room temperature. RNA was visualized by exposing the membrane to a phosphor screen followed by scanning the screen on a Typhoon 8600 scanner (Promega) in phosphor mode. Hybridization probe was made by PCR using oligonucleotides: 3Dseq-100-for (5’-GTT TGA AGG GGT GAA GGA A-3’) and 3Dseq 1085 rev (5’-CTC CCA TGT GAC TGT TTC AAA TG-3’) and pRLuc RA as a template. In the reaction, [α ^32^P] dATP (1 mCi/mL, 3000 Ci/mmol, GE Healthcare) was used with cold dNTPs (300 μM for each dCTP, dGTP, and dTTP and 10 μM for dATP) in total 100 μl reaction. The quality of PCR product was confirmed by agarose gel electrophoresis and cpm was determined by using scintillation counter (LKB Wallac 1217 Rackbeta liquid scintillation counter).

### One step growth curve assay

One day prior to infection, HeLa cells were seeded into 6-well plate at a density of 5 X 10^5^ cells/well. On the day of infection, the prepared HeLa monolayers were washed and infected with WT or EG virus at an MOI of 10 for 30 min at room temperature. Infected cells were washed once with PBS to remove unattached virus and then incubated at 37°C. At varying times post-infection, virus was harvested from cells and media by three freeze-thaw cycles with vortexing in between. The isolated virus was quantified by plaque assay.

### Virus quantification by plaque assay

For plaque assay, 0.5 X 10^6^ HeLa cells were plated one day prior to infection in 6-well plates. On the day of infection, the media was removed and the cells were washed once in PBS. Next, the cells were infected with the 10-fold dilutions of viral supernatants previously prepared and incubated for 30 min at 37°C. The inoculum was then removed and the cells were overlaid with 1% low-melting-point agarose (EMD). The overlay was allowed to solidify for 20 min at room temperature, and the plates were incubated at 37°C for 2 days to allow for plaque formation. 30 min prior to harvesting, the overlays were incubated with 4% formaldehyde/PBS to fix virus and cells. The agarose monolayers were then removed and plaques stained with crystal violet and counted. Viral titers were calculated in PFU/ml.

### Sorting subgenomic replicon expressing cells with flow cytometer

WT and GG-subgenomic replicon RNAs (2 μg of *in vitro* transcribed pRLuc RNA and 4 μg of *in vitro* transcribed pREGFP RNA) were electroporated into 1.2 X 10^7^ HeLa cells. The cells were incubated in normal growth media at 34°C and luciferase activity was monitored. When the specific activity of luciferase was about 10,000 RLU/μg of total protein (5 h post-transfection for pRLuc-/pREGFP-WT and 14 h post-transfection for pRLuc-/pREGFP GG), cells were harvested, chilled on ice for 5 min and sorted to collect transfection positive cells. For the sorting, EGFP positive cells were excited with a 488 Argon laser, detected with a 532/40 nm band pass filter and sorted by Cytopeia Influx sorter (Beckton-Dickinson, San Jose, CA) with 100 micron tip at 15 psi. The sorted HeLa cells were processed for transmission electron microscopy.

### Transmission electron microscopy

Virus-Infected HeLa cells or EGFP-positive post-sorted HeLa cells were harvested, fixed and embedded for TEM studies as described previously [43]. Briefly, the harvested cells were fixed with 1% glutaraldehyde, washed with 0.1 M cacodylate (Sodium dimethyl arsenate, Electron Microscopy Sciences) twice for 5 min each, incubated in 1% reduced osmium tetroxide containing 1% potassium ferricyanide in 0.1 M cacodylate for 60 min in the dark with one exchange and washed two times with 0.1 M cacodylate again. En bloc staining was performed with 3% uranyl acetate in 50% ethanol for 60 min in the dark. Dehydration was carried out with different concentrations of ethanol (50, 70, 95 and 100% for 5-10 min) and 100% acetonitrile. Embedding was performed overnight with 100% Epon at 65°C. The embedded sample was sectioned with a diamond knife (DiATOME) to slice 60-90 nm thickness by using ultra microtome (Reichart-Jung). The sectioned sample was place on copper grid (Electron Microscopy Science), stained with 2% uranyl acetate in 50% ethanol followed by lead citrate staining for 12 min. The grid was washed with H_2_O and the grid was dried completely. The image was obtained by using Jeol JEM 1200 EXII and FEI Tecnai G2 Spirit BioTwin located in Electron Microscopy Facility in Pennsylvania State University.

### Immunofluorescence

2.5 X 10^5^ cells were seeded on coverslips in 6-well plates and next day infected with WT or EG virus at an MOI of 10. GuHCl whenever used, the final concentration was 3 mM and the inhibitor was kept on cells with the virus for 30 min and for the entire incubation period thereafter. For subgenomic replicon RNA transfection, same number of cells were transfected with 2 μg pREGFP-GG RNA using the RNA transfection kit (Mirus). Infected or transfected cells were fixed with 4% formaldehyde at indicated times post-infection or transfection. Post fixation, cells were permeabilized with 20 μM digitonin for 10 min and washed with PBS. Cells were then blocked with 3% BSA in PBS for 1 h followed by incubation with anti-PI4P (1:200) antibody for 1 h in blocking buffer. This was followed by washing with PBS and incubating with Alexafluor594; IgM (1:1000) for 1 h. Finally, the cells were washed with PBS and mounted on slides using mounting buffer with DAPI. Samples were imaged with Zeiss Axiovert 200 M epifluorescence microscope.

## ACKNOWLEDGMENTS

We thank Dr. Greg Ning and staff of the electron microscopy core facility for their guidance and assistance.

